# Multispecies site occupancy modeling and study design for spatially replicated environmental DNA metabarcoding

**DOI:** 10.1101/2021.02.14.431192

**Authors:** Keiichi Fukaya, Natsuko Ito Kondo, Shin-Ichiro S. Matsuzaki, Taku Kadoya

## Abstract

1. Environmental DNA (eDNA) metabarcoding has become widely applied to gauge biodiversity in a noninvasive and cost-efficient manner. The detection of species using eDNA metabarcoding is, however, imperfect owing to various factors that can cause false negatives in the inherent multi-stage workflow.
2. Imperfect detection in the multi-stage workflow of eDNA metabarcoding also raises an issue of study design: namely, how available resources should be allocated among the different stages to optimize survey efficiency.
3. Here, we propose a variant of the multispecies site occupancy model for eDNA metabarcoding studies where samples are collected at multiple sites within a region of interest. This model describes the variation in sequence reads, the unique output of the high-throughput sequencers, in terms of the hierarchical workflow of eDNA metabarcoding and interspecific heterogeneity, allowing the decomposition of the sources of variation in the detectability of species throughout the different stages of the workflow. We also introduced a Bayesian decision analysis framework to identify the study design that optimizes the efficiency of species detection with a limited budget.
4. An application of the model to freshwater fish communities in the Lake Kasumigaura watershed, in Japan, highlighted a remarkable inhomogeneity in the detectability of species, indicating a potential risk of the biased detection of specific species. Species with lower site occupancy probabilities tended to be difficult to detect as they had lower capture probabilities and lower dominance of the sequences. The expected abundance of sequence reads was predicted to vary by up to 23.5 times between species.
5. An analysis of the study design suggested that ensuring multiple within-site replications of the environmental samples is preferred in order to achieve better species detection efficiency, provided that a throughput of tens of thousands of sequence reads was secured.
6. The proposed framework makes the application of eDNA metabarcoding more error-tolerant, allowing ecologists to monitor ecological communities more efficiently.

## 1. Introduction

High-throughput sequencing (HTS) technology has achieved a breakthrough in biology, particularly in the fields of ecology and evolution (Shokralla *et al*. 2012). In ecology, HTS can be applied alongside targeted PCR amplification to obtain the sequence data of target genetic markers (such as COI and rRNA gene regions that act as “DNA barcodes”) directly and effectively from environmental samples. This approach, called environmental DNA (eDNA) metabarcoding (Taberlet *et al*. 2012), enables the simultaneous detection of a number of species from environmental samples. Originally applied to profile microbial communities, the use of eDNA metabarcoding for macro-organisms is increasing and becoming a useful tool in ecology and conservation biology to gauge biodiversity in a noninvasive and cost-e*ffi*cient manner (Taberlet *et al*. 2012, Deiner *et al*. 2017).

The workflow of eDNA metabarcoding is multi-staged, requiring different procedures in the field and laboratory and on the computer (Deiner *et al*. 2017). First, in the field, researchers collect environmental samples from target sites to assess ecological communities. Then, the collected environmental samples are transferred to the laboratory where DNA molecules are extracted and amplified for the subsequent HTS of the target marker gene. Next, HTS yields a high number of raw sequence reads that are processed in a computer using bioinformatic tools, including quality filtering, read clustering, and assigning sequences to species (or, more generally, taxa) using a reference database. These eDNA metabarcoding procedures will eventually result in a species-by-sample table of sequence read counts, which can then be used to characterize communities in terms of the presence-absence (or even “relative abundance”) of species and measures of *α*-and *β*-diversity.

The detection of species using eDNA may be more sensitive than that of traditional survey methods (Smart *et al*. 2015, Yamamoto *et al*. 2017, McColl-Gausden *et al*. 2020). Nevertheless, varying detection probability can still lead to the imperfect detection of species, especially in community-wide eDNA metabarcoding, compared to that of targeted species-specific approaches such as quantitative PCR (Harper *et al*. 2018, Bylemans *et al*. 2019, Doi *et al*. 2019). False-negative detection errors are caused by a number of factors within the multi-stage eDNA metabarcoding procedure. For example, sampling processes, including both sample collection in the field and repeated subsampling for library preparation in the laboratory, can result in random losses of rare sequences (Zhan *et al*. 2014a, Shelton *et al*. 2016). Primer-template mismatches can have a huge impact on the e*ffi*ciency of PCR amplification and reduce the detection probability of species with low primer specificity (Piñol *et al*. 2015, Shelton *et al*. 2016). A smaller throughput (i.e., the total number of sequence reads) can lead to failures in the detection of rare sequences in the sample (Shelton *et al*. 2016). Sequence filtering procedures, used to remove artifactual sequences, may erroneously filter out actual target sequences (Zhan *et al*. 2014b). Some species can never be detected when the reference database is incomplete. Hence, for the reliable assessment of ecological communities using eDNA metabarcoding, it is crucial to adequately account for such detection errors that occur throughout the multi-stage procedure.

Quantitative knowledge of imperfect detection in eDNA metabarcoding can have important implications from the perspective of study design. To ensure the accurate assessment of species diversity using eDNA metabarcoding, researchers may choose more general sampling strategies, including increasing the number of sampling sites, increasing the number of replicates per site, or increasing the throughput. Given that budget is often limited, decisions must be made to allocate the available resources among these options. A key consideration when identifying optimal resource allocation is to evaluate the detectability of species at different stages of the eDNA metabarcoding workflow. This approach allows the prediction of the e*ffi*ciency of species detection among possible resource allocations under a given budget, which can then be used to determine the preferred study design.

To address these issues, we developed a novel extension of the multispecies site occupancy modeling framework (Dorazio & Royle 2005, Dorazio *et al*. 2006). Multispecies site occupancy models account for the imperfect detection of species in a metacommunity and have recently been applied in eDNA metabarcoding (Doi *et al*. 2019, Bush *et al*. 2020, McClenaghan *et al*. 2020, McColl-Gausden *et al*. 2020). However, to apply these models to eDNA metabarcoding data, the original sequence read counts must be aggregated into binary detection and non-detection indicators. We propose the adoption of an observation submodel based on a multinomial distribution, instead of the traditional Bernoulli observation submodel, to apply the multispecies site occupancy modeling framework directly to eDNA metabarcoding data. The objective of this approach is to allow the explicit decomposition of the sources of variation in the detectability of species in different stages of the process (Fig. 1). Furthermore, we introduce a Bayesian framework for optimizing resource allocation in a multi-stage workflow with a limited budget, which can be derived naturally based on the proposed model. We apply the proposed methods to an eDNA metabarcoding study of freshwater fish communities in the Lake Kasumigaura watershed, in Japan.

**Fig. 1.**
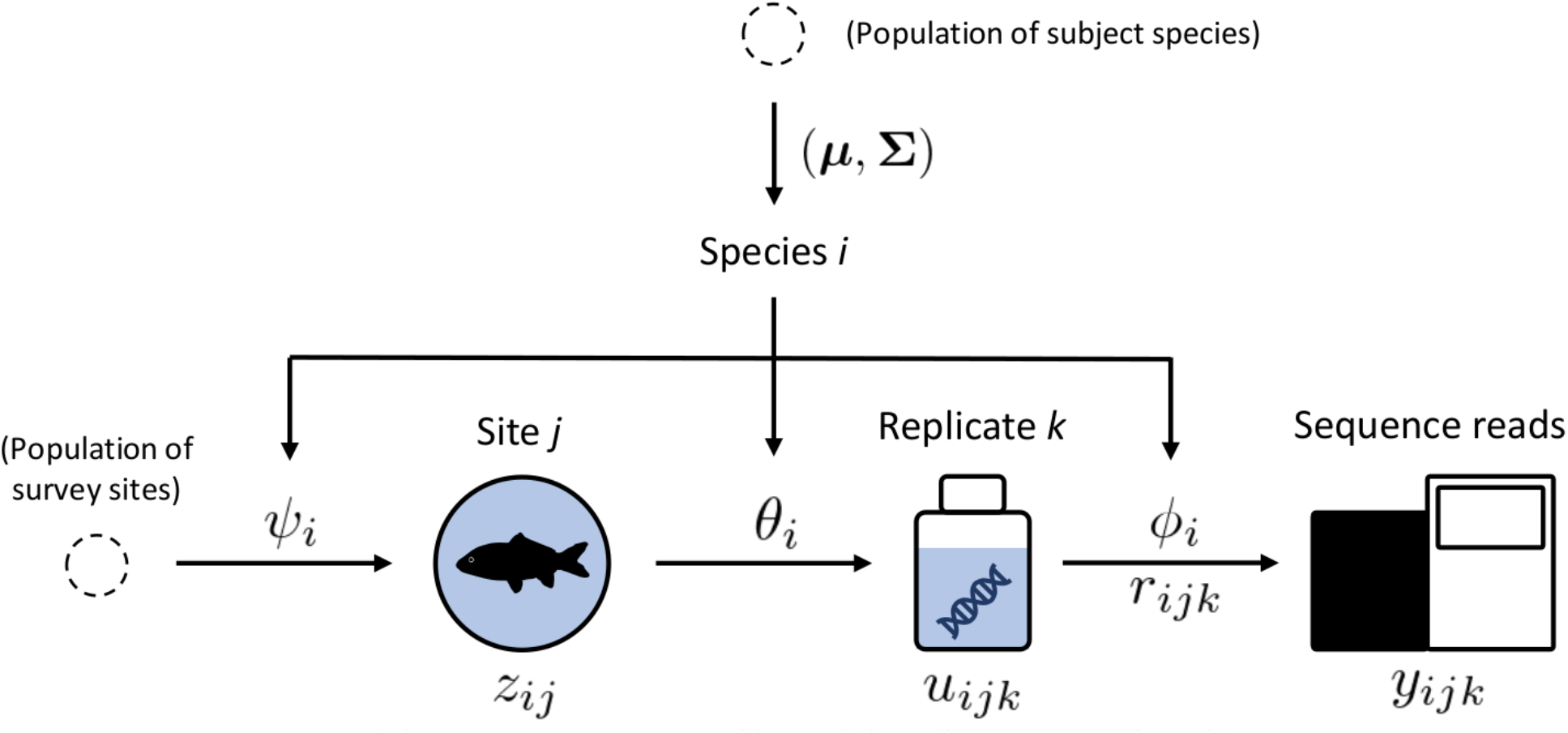
A diagram of the multispecies site occupancy model for spatially-replicated eDNA metabarcoding. A hierarchical dependence structure, characterized by a set of species-level parameters (occupancy probability *Ψ*, sequence capture probability *θ*, and relative dominance of sequence *ϕ*), is assumed between the latent variables *z* (site occupancy of species), *u* (presence or absence of DNA sequences of species in the replicate samples), and data *y* (sequence read counts). The sequence reads are assumed to follow a multinomial distribution that is conditional on the presence or absence (*u*) and relative dominance (*r*) of the sequences of all species, in addition to throughput. Species-level parameters are assumed to follow a community-level normal prior distribution with a mean vector ***µ*** and a covariance matrix **Σ**.

## 2 Hierarchical modeling for eDNA metabarcoding

### 2.1 Formulation

We considered studies of eDNA metabarcoding with spatially-replicated sampling designs (Fig. 1). Specifically, we assumed that the occurrence of *I* focal species was monitored at *J* sites sampled from an area of interest. We simply referred to taxonomic groups that emerged in an application of eDNA metabarcoding as “ species”. At site *j, K*_*j*_ replicates of environmental samples were collected. For each replicate, a unique sequence library was prepared to obtain separate sequence reads for each replicate. We denoted the resulting sequence read counts of species for replicate *k* (= 1, *…, K*_*j*_) at site *j* (= 1, *…, J*), obtained using HTS and subsequent bioinformatic processing, as ***y***_*jk*_ = (*y*_1*jk*_, *…, y*_*Ijk*_). We denoted the per-sample total sequence reads, which we called “throughput”, as *N*_*jk*_ =∑_*i*_ *y*_*ijk*_. Furthermore, we described a basic model structure with the least number of parameters; this model can be readily extended to account for additional heterogeneity, as described in Section 2.3.

We assumed that, conditional on the throughput *N*_*jk*_, sequence reads ***y***_*jk*_ followed a multinomial distribution (Shelton *et al*. 2016, Harrison *et al*. 2020). For *j* and *k* satisfying *N*_*jk*_ *>* 0, the observation model for ***y***_*jk*_ is expressed as

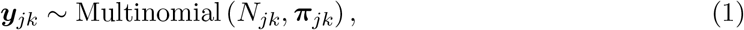

where ***π***_*jk*_ = (*π*_1*jk*_, *…, π*_*Ijk*_) are the multinomial cell probabilities expressing the relative frequency of the sequence of each species in the library of replicate *k* at site *j*. This observation model defines the per-sample marginal probability of species detection as 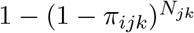, implying that species detection through HTS depends on both the relative frequency of the species sequence and the throughput. We model *π*_*ijk*_ as follows:

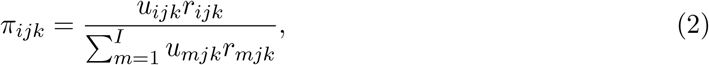

where *u*_*ijk*_ is a latent indicator variable representing the inclusion of the sequence of species *I* in the library of replicate *k* at site *j* and *r*_*ijk*_ is a latent variable that is proportional to the relative frequency of the sequence of species *i*, conditional on its presence in the library of replicate *k* at site *j*. Therefore, we decomposed the relative frequency of the sequence of species into binary components defining the presence or absence of the sequence in the library (*u*) and quantitative components defining the degree of dominance of the sequence (*r*). The expected frequency of species *i* in the sequence reads of replicate *k* at site *j* was assumed to be proportional to *r*_*ijk*_ when the library included the sequence of species (*u*_*ijk*_ = 1), whereas it was assumed to be zero when the library did not contain the sequence of species (*u*_*ijk*_ = 0). We noted that Equation 2 requires 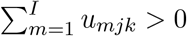; this is assured by defining the observation model (Equation 1) as conditional on a positive throughput (*N*_*jk*_ *>* 0), which implies that *u*_*ijk*_ is positive for one or more species.

The inclusion of the sequence of species can be modeled with a Bernoulli distribution as follows:

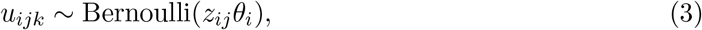

where *z*_*ij*_ is a latent indicator variable representing the occupancy of species *i* at site *j* and *θ*_*i*_ is the probability of per-replicate sequence capture that is conditional on the site occupancy of species *i*. Equation 3 implies that a replicate library includes the sequence of species *i* with probability *θ*_*i*_ when the species occupies site *j* (*z*_*ij*_ = 1), whereas replicate libraries of unoccupied sites (*z*_*ij*_ = 0) never contain the sequence of the species. In practice, *θ*_*i*_ could depend on the concentration of sequence copies in the field and the volume of collected medium.

The latent variable *r*_*ijk*_ is assumed to follow a gamma distribution as follows:

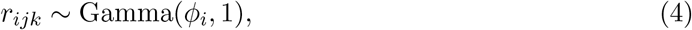

where we denote a gamma distribution with shape *a* and rate *b* by Gamma(*a, b*). Then, ***π***_*jk*_ follows a zero-inflated version of the Dirichlet distribution (Tang & Chen 2019), with a vector of expected values conditional on (*u*_1*jk*_, *…, u*_*Ijk*_) of (*u*_1*jk*_*ϕ*_1_*/A*_*jk*_, *…, u*_*Ijk*_*ϕ*_*I*_*/A*_*jk*_), where *A*_*jk*_ = ∑_*m*_ *u*_*mjk*_*ϕ*_*m*_. Therefore, *ϕ*_*i*_ is a species-specific parameter that governs the relative dominance of the sequence. Species that tend to yield more sequence reads, when they are present, are characterized by larger values of *ϕ*_*i*_. The relative dominance of sequences may vary depending on the relative abundance of species and the relative efficiency of PCR amplification.

We assume that the latent indicator of site occupancy *z*_*ij*_ follows a Bernoulli distribution as follows:

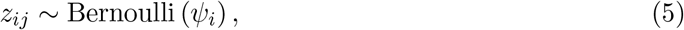

where *ψ*_*i*_ represents the site occupancy probability of species *i*. We noted that, in the context of eDNA-based monitoring, the site occupancy of species should be defined by the presence of DNA sequences rather than the presence of individuals in the site, as the presence of species can only be confirmed by the detection of sequences. In a lotic environment, such as rivers, long-distance transport of eDNA may result in the presence of sequences at sites where no individuals exist in the vicinity (Nukazawa *et al*. 2018). Consequently, general care should be taken when interpreting site occupancy when it is inferred from eDNA.

The model is characterized by a set of species-level parameters (*ϕ*_*i*_, *θ*_*i*_, *ψ*_*i*_). We can regard these parameters as random effects and assume that they follow a prior community-level distribution as follows (Dorazio & Royle 2005, Dorazio *et al*. 2006):

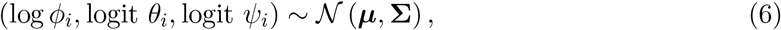

where ***µ*** and **∑** are the community mean vector and the covariance matrix of (*ϕ*_*i*_, *θ*_*i*_, *ψ*_*i*_) on the appropriate link scales. This formula allows information to be shared between species, thereby facilitating parameter inferences for the entire community, including for rare species (Iknayan *et al*. 2014).

A key assumption of the model is the absence of false positives; however, they can still occur in eDNA metabarcoding. We discuss a general consideration for the issue of false positives in Section 5.

### 2.2 Inference

With a fully Bayesian approach using Markov chain Monte Carlo (MCMC) algorithms, we can fit the model and conduct flexible posterior analyses. For this approach, we needed to specify prior distributions for the model parameters (***µ*, ∑**) in order to obtain samples from the posterior of all unknown quantities (*r, u, z, ϕ, θ, ψ*, ***µ*, ∑**). Specifically, for the mean vector ***µ***, we can specify a vague normal prior for each element. Furthermore, for the covariance matrix **∑**, we can specify a vague prior for each of the scale and correlation elements (Dorazio & Royle 2005, Dorazio *et al*. 2006). Posterior samples can be readily obtained using standard general-purpose MCMC software, for example JAGS (Plummer 2003; see the following application).

The process of species detection using eDNA metabarcoding can be characterized based on the resulting posterior of parameters. Specifically, the characteristics of individual species detection at different stages (Fig. 1) are represented by the posterior distribution of the species-level parameters *p*(*ϕ*_*i*_ | ***y***), *p*(*θ*_*i*_ | ***y***), and *p*(*ψ*_*i*_ | ***y***). In contrast, community-level characteristics can be inferred based on the posterior of the parameters in the community-level priors *p*(***µ*** | ***y***) and *p*(**∑** | ***y***), which represent the community mean and interspecific variation in the species-level parameters, respectively.

The occupancy of species at particular sites can be inferred based on the posterior probability Pr(*z*_*ij*_ = 1 | ***y***). Under the assumption that there are no false positives, the obvious solution Pr(*z*_*ij*_ = 1 | ***y***) = 1 is obtained in cases where the sequence of species *i* is detected at site *j*. In cases of non-detection, the posterior probability Pr(*z*_*ij*_ = 1 | ***y***) is less than one and can be evaluated based on the posterior samples of *z*_*ij*_. We can also infer the number of species that occur at a particular site, which can be expressed as ∑_*i*_ *z*_*ij*_ (Dorazio & Royle 2005) as a derived quantity.

### 2.3 Covariates and random effects for species-level parameters

Covariates and random effects can be modeled as the linear predictors of species-level parameters on appropriate link scales. The addition of such effects allows us to account for the heterogeneities in species-level parameters in a flexible manner. Covariates and random effects can be species-specific, site-specific, or replicate-specific. Possible covariates include site-specific environmental variables for site occupancy probability *ψ*, replicate-specific volume of filtered water for sequence capture probability *θ*, and a species-specific degree of primer-template mismatches for the relative dominance of sequences *ϕ*; see Section 4 for an illustration. Note that for models with such additional terms, the prior specification described in Section 2.2 will no longer apply. Depending on the parameters included, appropriate priors need to be specified.

## 3 Analysis for optimal study design

### 3.1 An overview

The problem of identifying the optimal study design for site occupancy studies was formulated by the criterion of maximizing the accuracy of the occupancy estimator under a given amount of survey budget or effort (or minimizing the amount of survey budget to achieve the desired estimator accuracy). This approach may be readily applicable to the simplest single-species site occupancy models with a large sample size, for which an analytical expression for the variance of the estimator is available (MacKenzie & Royle 2005, Guillera-Arroita *et al*. 2010, 2014, Lugg *et al*. 2018). A numerical simulation approach can be used for other classes of site occupancy models (Bailey *et al*. 2007); however, this may require extensive computation resources, especially for multispecies models (Sanderlin *et al*. 2014).

Here, as an alternative approach that is less computationally expensive, but still useful, we introduced a Bayesian decision analysis framework to determine the optimal study design under uncertainty (Chaloner & Verdinelli 1995, Dorazio & Johnson 2003, Guillera-Arroita *et al*. 2014). In this framework, the optimal study design is identified based on a profile of values of some arbitrary utility function averaged over the posterior (or even prior) distribution of unknown quantities. As a utility function relevant to eDNA metabarcoding, we considered the efficiency of species detection which can be derived from the multispecies site occupancy model described in Section 2.1.

We considered two different situations for optimizing study design under a given budget: to maximize survey efficiency for the assessment of (1) local species diversity and (2) regional species diversity. In the former, we infer the situation of balancing the number of replicates and the amount of throughput for given sites, to maximize the opportunity of detecting the occupying species at each site. In the latter, we infer the situation of balancing the number of sites, number of replicates, and the amount of throughput to maximize the number of species in the region expected to be detected. In the following, we describe a method for the former situation in detail. The method can be extended to apply to the latter situation, which we describe in detail in Appendix S1.

### 3.2 Study design for local species diversity

To model local species diversity, we assume that the occurrence of focal species *I* on sites *J* is modeled as described in Section 2.1. We used the result of model fitting, namely the posterior distribution, to obtain a profile of the efficiency of species detection among different resource allocations between the number of replicates per site and per throughput. The posterior distribution can be replaced by a prior distribution that reflects some guesses about the value of unknown quantities when no data was available at the beginning of the study (see Section 5). We briefly denoted the posterior distribution as *p*(***ξ*** | ***y***) by expressing all unknowns collectively as ***ξ***.

First, a cost function is introduced. As the number of sites is fixed, we need to consider only the costs associated with the procedures required after the collection of the environmental samples at each site. Specifically, we assumed the costs of HTS and library preparation (including filtration, DNA extraction, and PCR). We denoted the cost per sequence read for HTS as *λ*_1_ and the cost per replicate for library preparation as *λ*_2_. If we assume the same number of replicates per site and a constant throughput, the total cost for a study design with *J* sites, *K* replicates, and throughput *N* is expressed as *λ*_1_*JKN* + *λ*_2_*JK*. The total cost must be less than or equal to a given amount of budget *B*. The budget constraint can be expressed as:

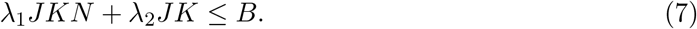

We assumed that we can obtain the maximum number of sequence reads per replicate, *N* = (*B* − *λ*_2_*JK*)*/λ*_1_*JK*, for *K* = 1, 2, *…*, ⌊*B/λ*_2_*J*⌋, where ⌊·⌋ represents the floor function.

Next, we defined utility as the expected number of detected species per site. This criterion is legitimate as it can be used to maximize the opportunity to verify the site occupancy of species at the level of local communities. Under the model described in Section 2.1, the expected number of detected species per site conditional on ***r*** = {*r*_*ijk*_} and ***u*** = {*u*_*ijk*_}, denoted as *U* (*K* | ***r, u***), is expressed as:

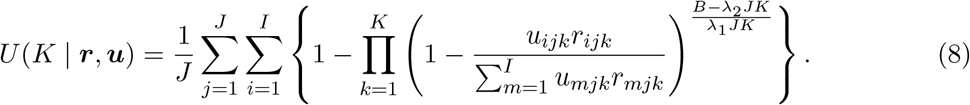

for *K* = 1, 2, *…*, ⌊*B/λ*_2_*J*⌋. By taking the average of *U* (*K* | ***r, u***) over the posterior predictive distribution of ***r*** and ***u***, we obtain the expected utility *U* (*K*) as follows:

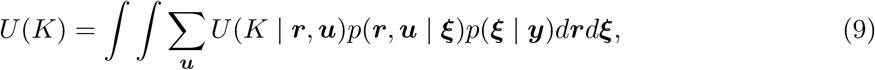

where *p*(***r, u*** | ***ξ***) represents the joint distribution of ***r*** and ***u***, conditional on ***ξ***, as specified by Equations 3 and 4. Note that, as is evident from taking the average over the posterior distribution, *U* (*K*) fully accounts for uncertainty in the unknowns of the model. When a large number of posterior samples are available, Equation 9 can be evaluated using the Monte Carlo integration by plugging the posterior predictive samples of *u* and *r* into Equation 8 and taking the sample average.

The preferred value of *K* (and the corresponding value of *N*) can be identified based on the profile of *U* (*K*). Note, however, that in practical applications, the budget amount itself is often considered. In this case, we could obtain a profile of *U* (*K*) for each possible budget amount and compare it between the budget levels to guide decision making (see Section 4).

## 4 Applications in fish eDNA metabarcoding

### 4.1 Materials and Methods

We applied the proposed model to a study of freshwater fish communities conducted in the Lake Kasumigaura watershed, Japan. The details of data acquisition and model fitting are fully described in Appendix S2. In August 2017, we collected three environmental samples (i.e., replicates) at 50 sites across the watershed (Matsuzaki *et al*. 2019). We used the universal primer set MiFish-U (Miya *et al*. 2015) to amplify a short fragment of fish DNA from the samples and sequenced the amplified DNA fragments using an MiSeq sequencer (Illumina, San Diego, CA, USA). We detected 50 freshwater fish taxa (henceforth referred to as “species”) which are listed in Table S1. In the following analysis, we omitted six of the 150 samples as no sequence reads were obtained. The statistical distribution of the throughputs of the remaining samples was right-skewed (Fig. S1), with a mean of 77, 910 and a standard deviation of 98, 035.

We analyzed the dataset using the proposed multispecies site occupancy model, with the covariate effects of the degree of primer-template mismatches on *ϕ* and that of lack of vegetation on *ψ*. The covariate effects were modeled as follows:

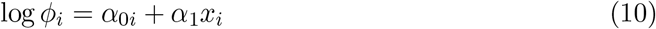

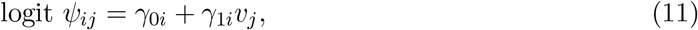

where *x*_*i*_ represents the degree of primer-template mismatches (total number of mismatched bases in the priming region of the forward and reverse primers; Table S1) of species *i*, and *v*_*j*_ indicates whether the riverbank at site *j* lacks aquatic and riparian vegetation. We treated *α*_0*i*_, *γ*_0*i*_, and *γ*_1*i*_ as random species effects. The community-level prior distribution for species-level parameters was then specified, as follows, instead of Equation 6:

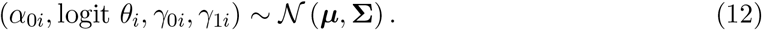

The model was fitted with JAGS software (Plummer 2003), in which we specified a vague prior for each parameter.

With the posterior samples, we examined profiles of the efficiency of species detection, as described in Section 3.2. We set *λ*_1_ = 0.01 JPY and *λ*_2_ = 5, 000 JPY based on approximative estimates of the cost of HTS and library preparation as of January 2019, respectively. We set 875,000 JPY as the baseline budget level, which approximated the actual budget for our study. Given that the maximum number of replicates for this budget level was three, we examined profiles of the efficiency of species detection under two hypothetical scenarios. First, we considered a situation where decisions were made within the range of budget levels under which the maximum number of replicates was fixed at three (scenario 1). Specifically, we compared profiles of the expected number of species detected over a range of budget levels around the baseline, *B* = {775 × 10^3^, 825 × 10^3^, 875 × 10^3^, 925 × 10^3^, 975 × 10^3^}. In the second scenario (scenario 2), we considered a situation where decisions were made within a larger range of budget levels with a different maximum number of replicates. We compared profiles of the expected number of species detected over a range of budget levels, *B* = {437.5 × 10^3^, 875 × 10^3^, 1, 750 × 10^3^, 2, 625 × 10^3^, 3, 500 × 10^3^}, which corresponds to 0.5, 1, 2, 3, and 4 times the baseline budget, respectively. The throughput allocated under each budget level is shown in Table 1. Note that under these budget levels, tens of thousands of sequence reads were secured for all conditions (minimum throughput: 16,667 reads).

**Table 1.** Values of throughput under two scenarios for the study design analysis of eDNA metabarcoding on freshwater fish communities in the Lake Kasumigaura watershed. The throughput is obtained as *N* = (*B* − *λ*_2_*JK*)*/λ*_1_*JK* for *K* = 1, 2, *…*, ⌊*B/λ*_2_*J*⌋ where we set *J* = 50 sites, *λ*_1_ = 0.01 JPY, and *λ*_2_ = 5, 000 JPY; see Sections 3.2 and 4.1. The unit of budget is 10^3^ JPY.

| Number of replicates<br>($K$) | Budget ($B$) | | | | | | | | | |
| --- | --- | --- | --- | --- | --- | --- | --- | --- | --- | --- |
|  | Scenario 1 |  |  |  |  | Scenario 2 |  |  |  |  |
|  | 775 | 825 | 875 | 925 | 975 | 437.5 | 875 | 1,750 | 2,625 | 3,500 |
| 1 | 1,050,000 | 1,150,000 | 1,250,000 | 1,350,000 | 1,450,000 | 375,000 | 1,250,000 | 3,000,000 | 4,750,000 | 6,500,000 |
| 2 | 275,000 | 325,000 | 375,000 | 425,000 | 475,000 | — | 375,000 | 1,250,000 | 2,125,000 | 3,000,000 |
| 3 | 16,667 | 50,000 | 83,333 | 116,667 | 150,000 | — | 83,333 | 666,667 | 1,250,000 | 1,833,333 |
| 4 | — | — | — | — | — | — | — | 375,000 | 812,500 | 1,250,000 |
| 5 | — | — | — | — | — | — | — | 200,000 | 550,000 | 900,000 |
| 6 | — | — | — | — | — | — | — | 83,333 | 375,000 | 666,667 |
| 7 | — | — | — | — | — | — | — | — | 250,000 | 500,000 |
| 8 | — | — | — | — | — | — | — | — | 156,250 | 375,000 |
| 9 | — | — | — | — | — | — | — | — | 83,333 | 277,778 |
| 10 | — | — | — | — | — | — | — | — | 25,000 | 200,000 |
| 11 | — | — | — | — | — | — | — | — | — | 136,364 |
| 12 | — | — | — | — | — | — | — | — | — | 83,333 |
| 13 | — | — | — | — | — | — | — | — | — | 38,462 |

## 4.2 Results

### 4.2.1 Species-level parameters

Site occupancy probabilities (*ψ*) varied considerably among species and showed a bimodal distribution, with more species tending to show occupancy probabilities close to one and others close to zero (Fig. 2A). Overall, the occupancy probability of species was reduced in the absence of vegetation (Fig. 2A, orange diamonds). A negative effect of lack of vegetation was identified at the community-level; the posterior median and the 95% highest posterior density interval (HPDI) of the community mean of *γ*_1_ were −1.039 and (−1.360, −0.748), respectively.

**Fig. 2.**
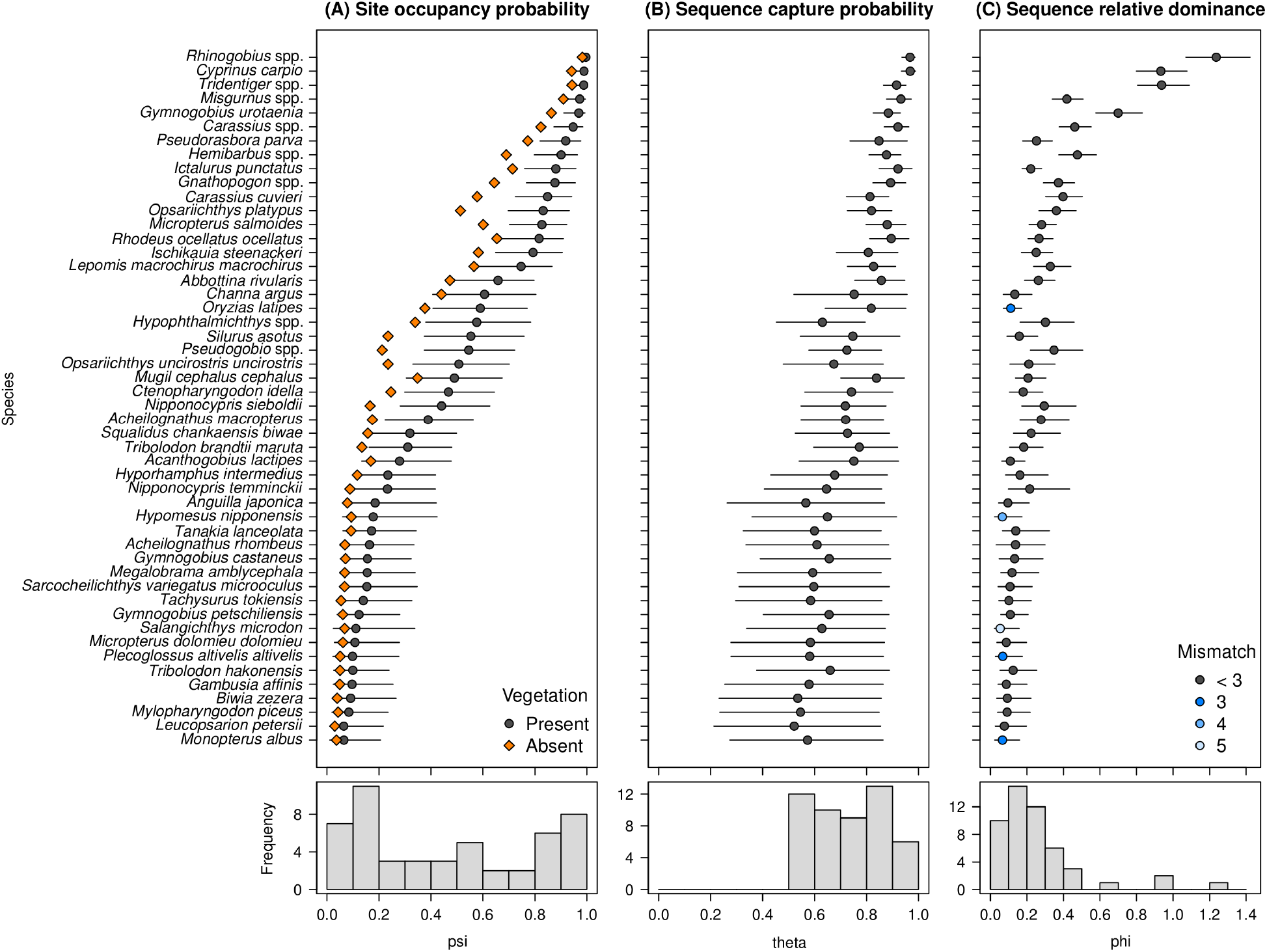
Estimates of species-level parameters: (A) site occupancy probabilities (*ψ*), (B) sequence capture probabilities (*θ*), and (C) sequence relative dominance (*ϕ*) for freshwater fishes in the Lake Kasumigaura watershed. In the upper panels, the posterior median and 95% HPDI for each species are represented by circles and lines, respectively. For site occupancy probability (A), presence (*v*_*j*_ = 0) and absence (*v*_*j*_ = 1) of vegetation at the site are distinguished by different symbols and colors. For the latter, we display only the posterior medians for visualization. For sequence relative dominance (C), the degree of primer-template mismatches is indicated by different colors. Species are listed in order of site occupancy probabilities. Lower panels show histograms of the posterior medians. For site occupancy probability (A), the histogram indicates the statistical distribution of posterior medians for vegetated sites.

Sequence capture probabilities (*θ*) also varied among species and tended to be uniformly distributed in the range of about 0.5 to 1 (Fig. 2B).

Sequence relative dominance (*ϕ*) showed a right-skewed distribution, with a small number of species exhibiting noticeably high values, whereas a large number of species showed low values (Fig. 2C). Species with a higher degree of primer-template mismatches tended to show lower sequence relative dominance (Fig. 2C, bluish circles). The posterior median of *α*_1_ was −0.195, whereas the 95% HPDI (−0.468, 0.115) overlapped with 0. The species ratio of sequence relative dominance predicted that, on average, and conditional on their presence, the species with the highest sequence dominance (*Rhinogobius* spp.) had 23.5 (95% HPDI; 5.7–54.3) times more sequence reads than the species with the lowest sequence dominance (*Salangichthys microdon*).

Estimates of the correlation coefficients between the species random effects are shown in Table 2. Positive correlations between the intercept of sequence relative dominance (*α*_0*i*_), the sequence capture probability (*θ*_*i*_), and the intercept of site occupancy probability (*γ*_0*i*_), were evident on the link scales. In contrast, there was no apparent correlation between the slope of site occupancy probability (*γ*_1*i*_) and other random effects.

**Table 2.** Posterior summary of correlation coefficients between the four species random effects. The posterior median and the 95 % HPDI (in brackets) are shown. Bold type indicates that the 95 % HPDI does not include zero.

| | $\alpha_{0i}$ | logit $\theta_i$ | $\gamma_{0i}$ | $\gamma_{1i}$ |
| --- | --- | --- | --- | --- |
| $\alpha_{0i}$ | — | — | — | — |
| logit $\theta_i$ | **0.62**<br>(0.17, 0.93) | — | — | — |
| $\gamma_{0i}$ | **0.82**<br>(0.65, 0.95) | **0.78**<br>(0.49, 0.97) | — | — |
| $\gamma_{1i}$ | −0.47<br>(−0.94, 0.14) | −0.21<br>(−0.78, 0.47) | −0.51<br>(−0.91, 0.16) | — |

### 4.2.2 Species site occupancy

The posterior probability of the site occupancy, Pr(*z*_*ij*_ = 1 | ***y***) of undetected species was lower than 0.15 in most cases (Fig. 3, bluish tiles), indicating that a reasonable effort had been made in species detection. Nevertheless, for some sites and species, we still found higher posterior probabilities of the site occupancy of undetected species (up to ∼ 0.3; Fig. 3, orangish tiles). The posterior probabilities of site occupancy tended to be higher in species with higher site occupancy probabilities (*ψ*) and lower sequence relative dominance (*ϕ*), such as *Channa argus* and *Oryzias latipes*.

**Fig. 3.**
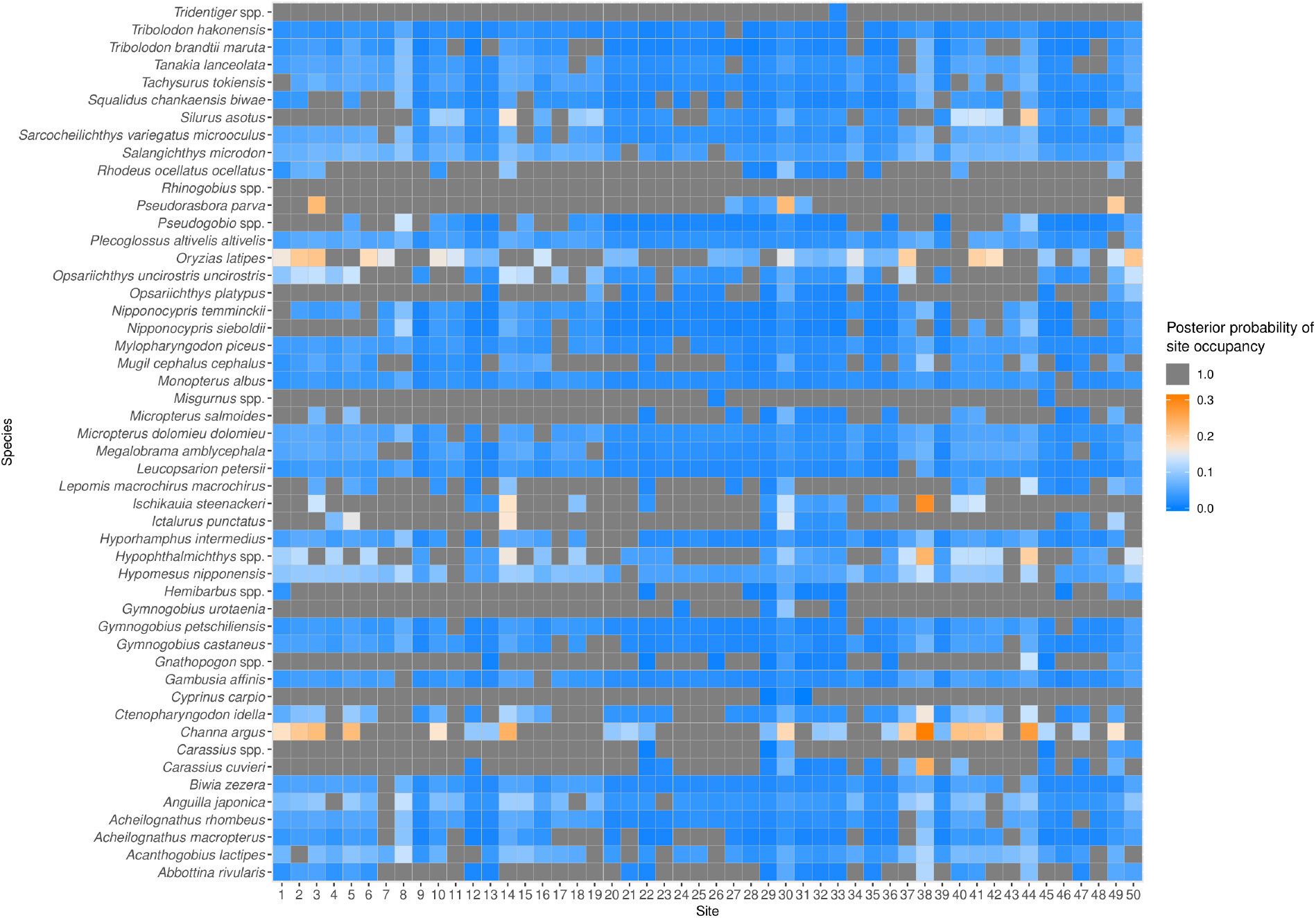
Tile plot of the posterior probabilities of species site occupancy. Gray tiles indicate that the species is detected; thus, the posterior probability is one.

#### 4.2.3 Study design analysis

In scenario 1 (Fig. 4A), the efficiency of species detection increased with the number of replicates, reaching a maximum at three replicates for all budget levels considered. At each number of replicates, the efficiency of species detection increased with the amount of budget, indicating that increased throughput (Table 1) also improves the efficiency of species detection. Nevertheless, under the range of budgets considered, the gain from increasing throughput was limited relative to increasing replicates.

**Fig. 4.**
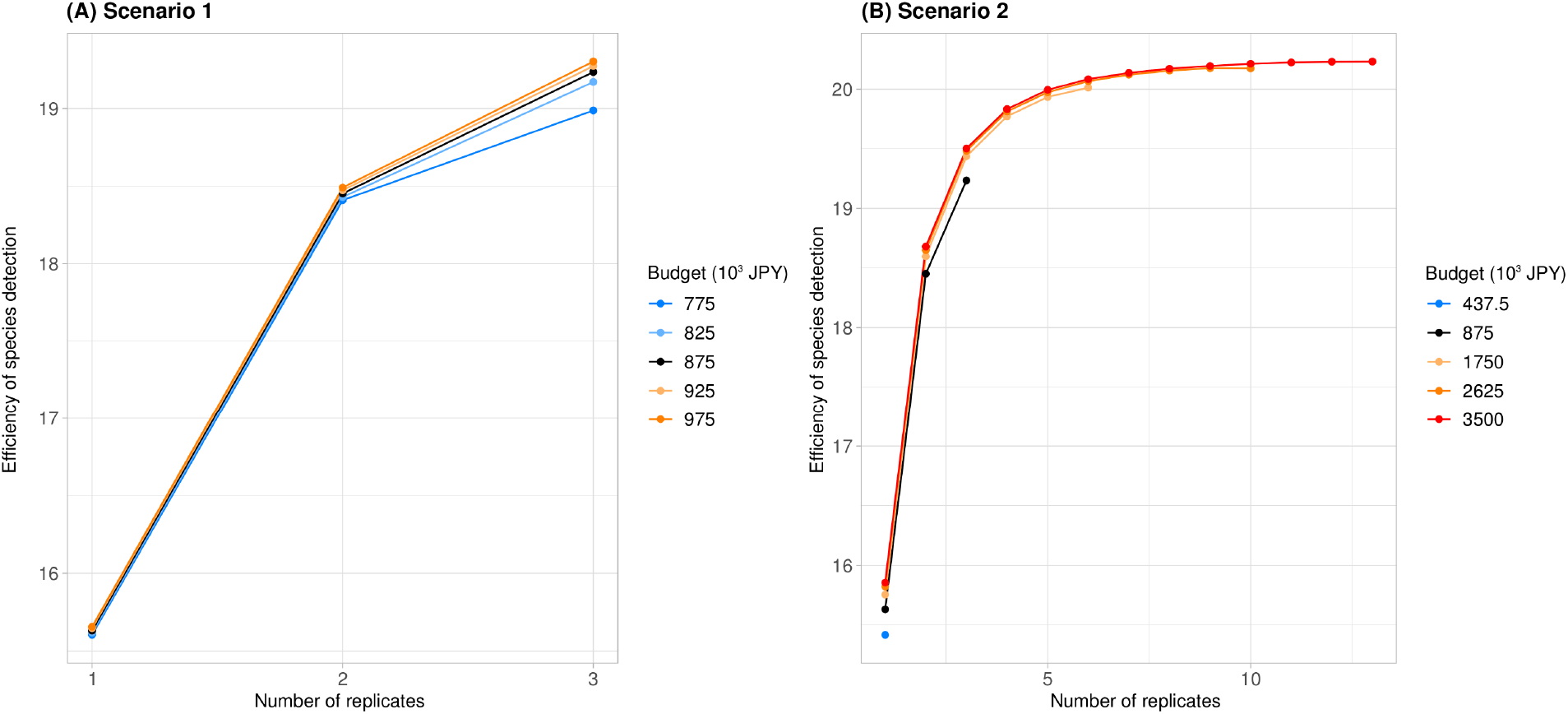
Profiles of the efficiency of species detection under specified amounts of research budgets for the eDNA metabarcoding of freshwater fish communities in the Lake Kasumigaura watershed. The efficiency of species detection is expressed by the expected number of species detected per site (see Section 3.2). The baseline profile is indicated by the black line. See Table 1 for the values of throughput for each condition.

In scenario 2 (Fig. 4B), the efficiency of species detection attainable varied significantly between budget levels. When the budget level was half of the baseline, only a single replicate could be taken and the efficiency of species detection was notably limited. When the budget level was several times greater than that of the baseline, it was possible to further improve the efficiency of species detection, which could be achieved by obtaining additional replicates. However, the magnitude of the gain in the efficiency of species detection rapidly decreased as the number of replicates increased. Thus, there are few additional benefits to spending more than three times the baseline budget.

## 5 Discussion

In this study, we proposed a novel variant of the multispecies site occupancy model (Dorazio & Royle 2005, Dorazio *et al*. 2006) for spatially-replicated eDNA metabarcoding that is directly applicable to sequence read counts, the unique output of recent HTS technology. With a series of submodels describing ecological and observation processes, the model describes the multi-stage process of species detection using eDNA metabarcoding and allows for the decomposition of the variability of sequence reads, in addition to the inference of site occupancy by species (Fig. 1). Furthermore, we introduced a Bayesian decision analysis framework of study designs for eDNA metabarcoding. These approaches are expected to make eDNA metabarcoding more efficient and robust for the imperfect detection of species.

In previous studies of eDNA metabarcoding, multispecies site occupancy models have been applied by aggregating sequence reads into binary detection/non-detection data (Doi *et al*. 2019, Bush *et al*. 2020, McClenaghan *et al*. 2020, McColl-Gausden *et al*. 2020). In such an approach, sequence read counts are not explicitly modeled, making it difficult to adequately decompose the combined effects of throughput and relative frequency of species sequences on the probability of species detection. By adopting a multinomial observation component, the proposed model overcomes this issue and thus provides a detailed description of the process of species detection using eDNA metabarcoding. The inference of the model is feasible with general-purpose software such as JAGS (Plummer 2003), as with the traditional multispecies site occupancy model (Dorazio & Royle 2005, Dorazio *et al*. 2006). The development of an R package is planned to make the methods introduced in this study even more convenient for researchers and practitioners to use.

An application to freshwater fish communities in the Lake Kasumigaura watershed highlighted the heterogeneity of the species-level parameters (Fig. 2). Covariate modeling revealed a tendency for species with a higher degree of primer-template mismatches to exhibit lower sequence relative dominance. Although the 95% HPDI of the coefficient for the effect of primer-template mismatch (*α*_1_) included zero, the observed tendency was consistent with previous results showing that primer-template mismatches can have a strong negative effect on the relative abundance of sequence reads owing to biased PCR amplification (Piñol *et al*. 2015, Shelton *et al*. 2016). The estimates of sequence relative dominance predict that there is a large inequality in the allocation of sequence reads among species; given that the majority of sequence reads are assigned to a small number of species, a false-negative detection error of a disproportionately large number of less dominant species is expected to occur at the stage of HTS when throughput is limited. In addition, the correlations of species random effects (Table 2) imply that rare species with lower site occupancy probability tend to be difficult to detect when they are present, compared to the detection of common species with higher occupancy probability. These results, revealed by the use of the proposed model, underscore the large inhomogeneity in eDNA metabarcoding, which can result in the biased detection of specific species.

Analysis of the study design suggested that multiple replicates can result in better efficiency in species detection. In the eDNA metabarcoding of freshwater fish communities in the Lake Kasumigaura watershed, the number of species detected per site was expected to increase with an increase in the number of replicates (Fig. 4), as long as a throughput of tens of thousands of sequence reads was secured (Table 1). In reality, obtaining a small number of replicates can also be an efficient strategy for assessing regional species diversity (Fig. S2). We note, however, that this finding may be specific to the application of the model to the freshwater fish communities in the Lake Kasumigaura watershed. It is likely that the optimal balance between throughput and the number of replicates (and even the number of sites) can vary depending on, for example, the distribution of species-level parameters, the costs required for different stages of the workflow, or the amount of research budget available. Therefore, a further examination into different systems and conditions is required.

In the Bayesian decision analysis framework for the study design, we used the posterior distribution *p*(***ξ*** | ***y***) to obtain the expected utility *U* (*K*) (Equation 9). This approach allowed for rational decisions based on the dataset obtained. In contrast, when a dataset is not available, a prior distribution *p*(***ξ***), instead of a posterior distribution *p*(***ξ*** | ***y***), can be used to obtain the expected utility (Chaloner & Verdinelli 1995, Guillera-Arroita *et al*. 2014). Such a treatment could be used to design preliminary research and develop a sequential design approach as addressed earlier (Guillera-Arroita *et al*. 2014).

In eDNA metabarcoding, false positives can occur because of various technical factors, including contamination, amplification or sequencing errors, or tag switching (Zhan & MacIsaac 2015, Furlan *et al*. 2020). As unmodeled false positives can cause a significant bias in the inference of site occupancy (Royle & Link 2006), adequate precautions must be taken in applying the proposed model to an eDNA metabarcoding study. We recommend that efforts be made to reduce as many false positives as possible by, for example, performing careful experimental manipulations, utilizing negative controls, conducting conservative quality filtering, and using clustering and de-noising approaches. Generally, reducing false positives can lead to an increase in false negatives (Zhan *et al*. 2014b, Zhan & MacIsaac 2015). It would be reasonable, however, to make more efforts to avoid generating false positives than false negatives during the process of data acquisition as the proposed model formally accounts for false negatives under the assumption of the absence of false positives. It should be noted that the model never corrects false negatives caused by incomplete reference databases. As a complementary approach, further research is warranted to extend the model to account for false positives in addition to false negatives.

## Acknowledgements

We are grateful to the following people for their support in the fieldwork and laboratory experiments: Ayato Kohzu, Mirai Watanabe, Kazuhiro Komatsu, Haruyo Yamaguchi, Haruko Ando, Koichi Shimotori, Sayuri Nakamura, Megumi Nakagawa, Megumi Yoshiba, Junko Hirama, Tomomi Shindo, Junko Machizawa, and Nobuyoshi Nakajima. We thank I. K. Shimatani for providing valuable feedback on the earlier version of this manuscript. Funding was provided by the Environmental Restoration and Conservation Agency (ERTDF, No. 4–1705) and the Japan Society for the Promotion of Science (KAKENHI, Nos 20H03010 and 20K06102). This research was supported by allocation of computing resources of the HPE SGI 8600 supercomputer from the Institute of Statistical Mathematics. The authors have no conflicts of interest to declare.

## Author Contributions

KF and TK conceived the study and designed the methodology; SSM designed and organized the fieldwork; NK conducted eDNA metabarcoding and organized the dataset; KF analyzed the dataset; KF and NK drafted the manuscript. All authors contributed critically to the drafts and gave final approval for publication.

## Data Availability Statement

The raw sequence reads will be archived in the DDBJ Sequence Read Archive. The dataset and codes will be archived in the Dryad Digital Repository upon acceptance.

## Notes

### Competing Interest Statement

The authors have declared no competing interest.

