## Supporting information for "Multispecies site occupancy modeling and study design for spatially replicated environmental DNA metabarcoding"

#### Contents

**Appendix S1. Study design for regional diversity** **2**

**Appendix S2. The eDNA metabarcoding of freshwater fish communities** **4**
Data acquisition . . . . . . . . . . . . . . . . . . . . . . . . . . . . . . . . . . . 4
Model fitting . . . . . . . . . . . . . . . . . . . . . . . . . . . . . . . . . . . . . 5

**Appendix S3. Supplementary tables and figures** **7**

#### Appendix S1. Study design for regional diversity

We considered a study design problem that balanced the number of sites $J$, number of replicates per site $K$, and throughput $N$ under a given amount of budget, in order to maximize the number of species in the region expected to be detected. We inferred a situation where $J$ new sites were sampled from the study region to detect $I$ species of interest using eDNA metabarcoding.

The cost function should constitute a new term on site visits. Although the cost of visiting a site may depend on the distance (Field *et al.* 2005), here, we simply assumed that the visiting cost per site was constant. We denoted the cost for HTS per sequence read as $\lambda_1$, the cost for library preparation per replicate as $\lambda_2$, and the visiting cost per site as $\lambda_3$. The budget constraint is expressed as:

$$\lambda_1 JKN + \lambda_2 JK + \lambda_3 J \leq B', \tag{S1}$$

where $B'$ is a given amount of budget [we used a dash ($'$) to distinguish it from $B$, the amount of budget for HTS and library preparation, in the main text]. We assumed that we could obtain the maximum number of sequence reads per replicate, $N = (B' - \lambda_2 JK - \lambda_3 J)/\lambda_1 JK$, for $J$ and $K$ satisfying $B' - \lambda_2 JK - \lambda_3 J > 0$.

We defined utility as the expected total number of species detected in $J$ sites. Under the model described in Section 2.1 in the main text, the conditional utility, denoted as $U(J, K \mid \boldsymbol{r}, \boldsymbol{u})$, is expressed as:

$$U(J, K \mid \boldsymbol{r}, \boldsymbol{u}) = \sum_{i=1}^{I} \left\{ 1 - \prod_{j=1}^{J} \prod_{k=1}^{K} \left( 1 - \frac{u_{ijk} r_{ijk}}{\sum_{m=1}^{I} u_{mjk} r_{mjk}} \right)^{\frac{B' - \lambda_2 JK - \lambda_3 J}{\lambda_1 JK}} \right\}, \tag{S2}$$

for $J$ and $K$ satisfying $B' - \lambda_2 JK - \lambda_3 J > 0$. By taking the average of Equation S2 over the posterior predictive distribution of $\boldsymbol{r}$ and $\boldsymbol{u}$, we obtained the expected utility $U(J, K)$ as follows:

$$U(J, K) = \int \int \sum_{\boldsymbol{u}, \boldsymbol{z}} U(J, K \mid \boldsymbol{r}, \boldsymbol{u}) p(\boldsymbol{r}, \boldsymbol{u} \mid \boldsymbol{z}, \boldsymbol{\xi}) p(\boldsymbol{z} \mid \boldsymbol{\xi}) p(\boldsymbol{\xi} \mid \boldsymbol{y}) d\boldsymbol{r} d\boldsymbol{\xi}, \tag{S3}$$

where $p(\boldsymbol{r}, \boldsymbol{u} \mid \boldsymbol{z}, \boldsymbol{\xi})$ and $p(\boldsymbol{z} \mid \boldsymbol{\xi})$ represents the joint distribution of $\boldsymbol{r}$ and $\boldsymbol{u}$ conditional on $\boldsymbol{z} = \{z_{ij}\}$ and $\boldsymbol{\xi}$ and the joint distribution of $\boldsymbol{z}$ conditional on $\boldsymbol{\xi}$, respectively. Equation S3 can be evaluated using the Monte Carlo integration. Specifically, for each MCMC replicate $m = 1, \ldots, M$, we sampled $r_{ijk}^{(m)}$ and $u_{ijk}^{(m)}$ according to the submodels $r_{ijk}^{(m)} \sim \text{Gamma}(\phi_i^{(m)}, 1)$, $u_{ijk}^{(m)} \sim \text{Bernoulli}(z_{ij}^{(m)} \theta_i^{(m)})$, and $z_{ij}^{(m)} \sim \text{Bernoulli}(\psi_i^{(m)})$, and then evaluated the sample mean of Equation S2.

When the heterogeneity of species-level parameters is modeled, as described in Section 2.3 in the main text, it can be accounted for in the evaluation of the expected utility. For this, we can sample the values of the covariates and/or the random effects from a parametric model representing their statistical distribution to obtain heterogeneous species-level parameters. It should be noted, however, that these additional sampling processes can require more Monte Carlo iterations to obtain precise estimates of expected utility.

We show the profile of $U(J, K)$ for $K = (1, 2, 3, 4, 8, 16)$ replicates in Fig. S2, obtained using the results from the analysis of the freshwater fish assemblages in the Lake Kasumigaura watershed. We set $\lambda_1 = 0.01$ JPY, $\lambda_2 = 5,000$ JPY, $\lambda_3 = 5,000$ JPY, and $B' = 1,125,000$ JPY. The value of $B'$ represents the approximate total budget for the study. It was obtained as $B' = B + \lambda_3 J$, where $B = 875,000$ JPY was the amount of budget for library preparation and HTS, and $J = 50$ was the number of sites surveyed in the study (see Section 4.1 in the main text). We evaluated $U(J, K)$ for sites with vegetation.

Fig. S2 indicates that, in general, surveying more sites increases the efficiency of species detection. However, due to a smaller throughput, detection efficiency can decrease when the number of sites becomes excessive. The attainable species detection efficiency was maximized when the number of replicates was two (detection of 48.70 species was expected with $J = 71$ sites and $N = 42,254$ sequence reads), followed by that obtained from three replicates (detection of 48.66 species was expected with $J = 52$ sites and $N = 54,487$ sequence reads) and four replicates (detection of 48.52 species was expected with $J = 43$ sites and $N = 29,070$ sequence reads). When the number of replicates was greater, however, the attainable species detection efficiency was considerably limited. This result suggests that, in the freshwater fish communities of interest, it is preferable to ensure a small number of replicates to evaluate regional species diversity.

### Appendix S2. The eDNA metabarcoding of freshwater fish communities

##### Data acquisition

We collected river water samples at 50 sites in the Lake Kasumigaura watershed over two days in August 2017. The general characteristics of the study area and the selection of the sites are presented in Matsuzaki *et al.* (2019). Although there was a small amount of precipitation (8.5 mm cumulative) in the study area two days prior to sampling, samples were collected under base-flow conditions (Matsuzaki *et al.* 2019).

At each site, we obtained three environmental samples (i.e., replicates) by collecting 1 L of river water at the center of the river and near the left and right riverbanks. The samples of river water were placed in hypochlorous acid-treated polypropylene containers, before immediately being stored on ice.

On the date of collection, we filtered the river water samples under negative pressure using 47-mm diameter Whatman GF/F glass microfiber filters (Cytiva, Tokyo, Japan). As the filters clogged easily, we used up to four filters to filtrate each 1-L sample of river water. We then wrapped the filters in aluminum foil and stored them at −28 °C until DNA extraction.

We extracted DNA from the filters using the DNeasy Blood & Tissue Kit (QIAGEN, Hilden, Germany) according to the manufacturer's protocol and Uchii *et al.* (2016). The reactions were performed with Buffer AL and Proteinase K for each filter. The spin column was then applied to the reaction solutions and eluted with 100 ml AE buffer to obtain a template for PCR. Where there were multiple filters per replicate, we mixed the reaction solutions of up to two filters when applying the spin column, and mixed equal amounts of eluted solutions obtained from the same sample.

We used the universal primer set MiFish-U (Miya *et al.* 2015), which amplifies a hypervariable region of the 12S rRNA gene. The total volume of the first PCR reaction was 12 µl, containing 2 µl of template DNA, 6 µl of 2x PCR Buffer, 0.3 µM of MiFish-U-F and -R primers with adapter sequences, 0.4 mM of dNTPs, and 0.24 U of KOD FX Neo (TOYOBO, Osaka, Japan). The cycling condition of the first PCR was 94 °C for 2 min of initial denaturing, followed by 35 cycles of 98 °C for 10 sec, 61 °C for 30 sec, 70 °C for 30 sec, and 5 min at 70 °C for the final extension. For samples that showed no or little amplification, we performed a touchdown PCR by decreasing the annealing temperature by 2 °C from 67 °C to 59 °C for 5 cycles each, then performing 15 cycles at 59 °C. The products of the first PCR were purified using AMPure XP (Beckman Coulter Inc., Brea, CA, USA) according to the manufacturer's instructions. We conducted the second PCR to add adapters and indices for sample identification with the same reaction solution composition as the first PCR under the following cycling conditions: 94 °C for 2 min of initial denaturing, followed by 25 cycles of 98 °C for 10 sec, 70 °C for 30 sec, and 5 min at 68 °C for the final extension.

We mixed an equal volume of the products of the second PCR of each replicate to obtain a sequence library. We purified the mixture using a QIAquick PCR Purification Kit (QIAGEN) and size-selected samples using E-Gel SizeSelect II (Thermo Fisher Scientific, Waltham, MA, USA). The concentration of the library was measured using a Qubit Fluorometer 3.0 (Thermo Fisher Scientific) with a Qubit dsDNA HS Assay Kit (Thermo Fisher Scientific) and diluted to 4 nM. After an alkaline denaturation, the library was sequenced in MiSeq (Illumina, San Diego, CA, USA) using the MiSeq Reagent Kit v2 for 300 cycles with 1% PhiX (Illumina).

The fastq files of the raw sequence reads was directly analyzed using the MiFish Pipeline on the MitoFish server (MiFish DB version 30; Sato *et al.* 2018) with all default settings. Briefly, the pipeline carried out filtering by quality, assembling paired-end reads, filtering by length (229 ± 25 bp), removing primer sequences, clustering at 99% identity, and blastn searching for representative sequences with more than 97% identity.

Taxonomic assignments of the representative sequences were performed up to the level of subspecies where possible. To eliminate apparent false positive detections and inaccurate taxonomic assignments, we reviewed the taxonomic assignments based on the blastn result, according to the procedure of Hayami *et al.* (2020), with some modification. Specifically, the taxonomic assignments were revised as follows: (1) we merged congeneric species that could not be distinguished based on the amplified region of 12S rRNA gene and assigned them to their genus. When it could be determined from existing literature that there was only a single taxon of the same genus in the study area, the sequences were reassigned to that taxon; (2) based on the existing literature, we omitted fish taxa that are assumed not to be present in the study area. However, when sequences were assigned to congeners of a taxon in the study area and there was only a single taxon of the same genus in the study area, they were reassigned to that taxon. We also omitted sequences of non-fish taxa.

In total, 50 freshwater fish taxa, including native, translocated, and exotic taxa, were detected and are listed in Table S1. In the model fitting, we omitted six of the 150 samples as no sequence reads were obtained. The statistical distribution of the throughput of the remaining samples was right-skewed (Fig. S1), with a mean of $77,910$ and a standard deviation of $98,035$.

##### Model fitting

We analyzed the dataset using the proposed multispecies site occupancy model, with the covariate effects of the degree of primer-template mismatches on $\phi$ and that of lack of vegetation on $\psi$. The covariate effects were modeled as follows:

$$\log \phi_i = \alpha_{0i} + \alpha_1 x_i \tag{S4}$$

$$\text{logit } \psi_{ij} = \gamma_{0i} + \gamma_{1i} v_j, \tag{S5}$$

where $x_i$ represents the degree of primer-template mismatches of species $i$ and $v_j$ indicates whether the riverbank at site $j$ lacks vegetation. For $x_i$, we used the total number of mismatched bases in the priming region of the forward and reverse primers (Piñol *et al.* 2015) which was identified for each species based on the mitochondrial genome sequences in the MitoFish database (version 3.63; Iwasaki *et al.* 2013). For species whose genome sequences were missing, the degree of mismatches for species of the same or most closely related genus was determined and those values were used instead (see Table S1 for details). For species whose genome sequence was missing, we searched for sequences of the mitochondrial 12S rRNA gene in the NCBI Nucleotide Database to identify their degree of mismatches. For species whose priming region sequence was still unknown, the degree of mismatches for species of the same or most closely related genus was determined, and those values were used instead (see Table S1 for details). For $v_j$, a binary variable was used that took the value of 1 if no aquatic and riparian vegetation existed at site $j$ because both sides of the river bank were straightened with concrete, and 0 otherwise. We identified the value of $v_i$ by visually checking approximately 50 m around the sampling locations for each site.

We treated $\alpha_{0i}$, $\gamma_{0i}$, and $\gamma_{1i}$ as species random effects. The community-level prior distribution for species-level parameters was then specified as follows:

$$(\alpha_{0i}, \text{logit } \theta_i, \gamma_{0i}, \gamma_{1i}) \sim \mathcal{N}\left(\boldsymbol{\mu}, \boldsymbol{\Sigma}\right). \tag{S6}$$

The model was fitted with JAGS software (Plummer 2003) in which we specified a prior distribution $\mathcal{N}(0, 10^3)$ for each element of $\boldsymbol{\mu}$ and $\alpha_1$, and Uniform$(0, 10^3)$ and Uniform$(-1, 1)$ for the scale and correlation element of the covariance matrix $\boldsymbol{\Sigma}$, respectively. We ran six independent Markov chains, each with 30,000 burn-in periods followed by 500,000 iterations and were thinned at intervals of 500 to obtain the posterior samples. We assessed the convergence of the posterior with the $\hat{R}$ statistic. The convergence was achieved at the recommended level ($\hat{R} < 1.1$) for all except a very few of the parameters of interest. The goodness-of-fit of the model was assessed using the Freeman–Tukey discrepancy statistic (Kéry & Royle 2016), which indicated no clear lack of model fit (Bayesian $p$-value: 0.82).

### Appendix S3. Supplementary tables and figures

Table S1. List of freshwater fish taxa detected at 50 sites in the Lake Kasumigaura watershed using eDNA metabarcoding with the MiFish-U primer set. The degree of primer-template mismatches is expressed as the total number of base mismatches in the binding region of the forward and reverse primers.

| Order | Family | Species | Degree of primer-template mismatches |
|---|---|---|---|
| Anguilliformes | Anguillidae | *Anguilla japonica* | 1 |
| Cypriniformes | Cyprinidae | *Abbottina rivularis* | 1 |
| | | *Acheilognathus macropterus* | 1 |
| | | *Acheilognathus rhombeus* | 1 |
| | | *Biwia zezera* | 1 |
| | | *Carassius cuvieri* | 1 |
| | | *Carassius* spp. | 1[a] |
| | | *Ctenopharyngodon idella* | 1 |
| | | *Cyprinus carpio* | 2 |
| | | *Gnathopogon* spp. | 1[b] |
| | | *Hemibarbus* spp. | 1[c] |
| | | *Hypophthalmichthys* spp. | 1[d] |
| | | *Ischikauia steenackeri* | 1 |
| | | *Megalobrama amblycephala* | 1 |
| | | *Mylopharyngodon piceus* | 1 |
| | | *Nipponocypris sieboldii* | 1 |
| | | *Nipponocypris temminckii* | 1 |
| | | *Opsariichthys platypus* | 1 |
| | | *Opsariichthys uncirostris uncirostris* | 1[e] |
| | | *Pseudogobio* spp. | 1[f] |
| | | *Pseudorasbora parva* | 1 |
| | | *Rhodeus ocellatus ocellatus* | 1[g] |
| | | *Sarcocheilichthys variegatus microoculus* | 1 |
| | | *Squalidus chankaensis biwae* | 1[h] |
| | | *Tanakia lanceolata* | 1 |
| | | *Tribolodon brandtii maruta* | 1[i] |
| | | *Tribolodon hakonensis* | 1 |
| | Cobitidae | *Misgurnus* spp. | 1[j] |
| Siluriformes | Siluridae | *Silurus asotus* | 1 |
| | Bagridae | *Tachysurus tokiensis* | 2 |
| | Ictaluridae | *Ictalurus punctatus* | 2 |
| Osmeriformes | Osmeridae | *Hypomesus nipponensis* | 4 |
| | Plecoglossidae | *Plecoglossus altivelis altivelis* | 3[k] |
| | Salangidae | *Salangichthys microdon* | 5 |
| Gobiiformes | Gobiidae | *Acanthogobius lactipes* | 2 |
| | | *Gymnogobius castaneus* | 1[l] |
| | | *Gymnogobius petschiliensis* | 1 |
| | | *Gymnogobius urotaenia* | 1 |
| | | *Leucopsarion petersii* | 2[m] |
| | | *Rhinogobius* spp. | 1[n] |
| | | *Tridentiger* spp. | 1[o] |
| Mugiliformes | Mugilidae | *Mugil cephalus cephalus* | 2[p] |
| Beloniformes | Adrianichthyidae | *Oryzias latipes* | 3 |
| | Hemiramphidae | *Hyporhamphus intermedius* | 1 |
| Cyprinodontiformes | Poeciliidae | *Gambusia affinis* | 2 |
| Synbranchiformes | Synbranchidae | *Monopterus albus* | 3 |
| Anabantiformes | Channidae | *Channa argus* | 0 |
| Perciformes | Centrarchidae | *Lepomis macrochirus macrochirus* | 1[q] |
| | | *Micropterus salmoides* | 1 |
| | | *Micropterus dolomieu dolomieu* | 1[r] |

[a] Used the most typical value in the same genus. [b] Used the value of *G. elongatus* present in the study area. [c] Used the value of *H. barbus* present in the study area. [d] Used the value of *H. molitrix* present in the study area. [e] Used the value of a species registered as *O. uncirostris*. [f] Used the value of *P. esocinus* present in the study area. [g] Used the value of a species registered as *R. ocellatus*. [h] Used the value of a species registered as *S. chankaensis*. [i] Used the value of a species registered as *T. brandtii*. [j] Used the value of *M. anguillicaudatus* present in the study area. [k] Used the value of a species registered as *P. altivelis*. [l] Used the same value of *G. petschiliensis* and *G. urotaenia*. [m] Used the value of a species (*Acanthogobius lactipes*) in the most closely related genus (Agorreta *et al.* 2013). [n] Used the most typical value in the same genus. [o] Used the value of *T. obscurus* present in the study area. [p] Used the value of a species registered as *M. cephalus*. [q] Used the value of a species registered as *L. macrochirus*. [r] Used the value of a species registered as *M. dolomieu*.

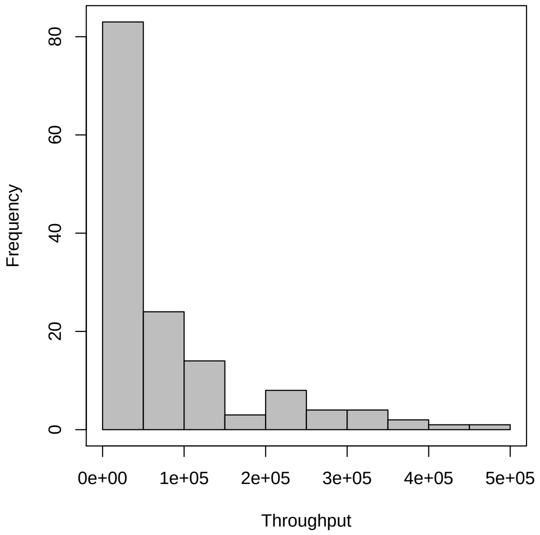


Fig. S1. Statistical distribution of throughput in the eDNA metabarcoding study of freshwater fish communities in the Lake Kasumigaura watershed.

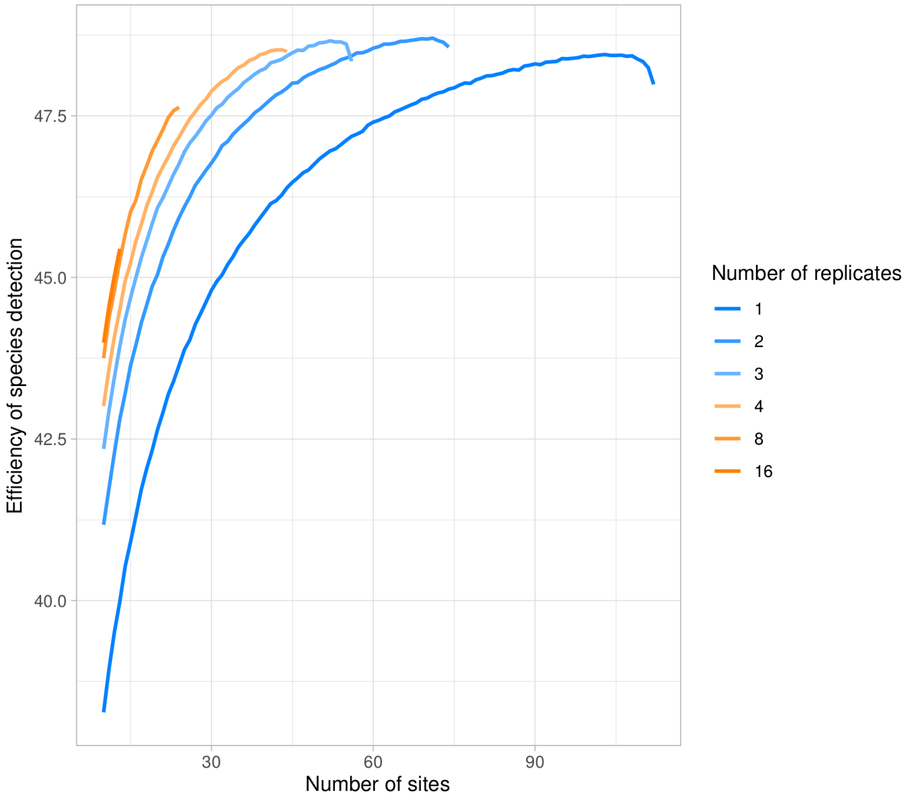


Fig. S2. The efficiency of species detection, represented by the number of species in the study region expected to be detected, expressed as a function of the number of sites and replicates. The profiles were obtained under specified amounts of research budget for the eDNA metabarcoding of freshwater fish communities in the Lake Kasumigaura watershed; see Appendix S1 for details.
